# Nanoscale Dynamics of Centromere Nucleosomes and the Critical Roles of CENP-A

**DOI:** 10.1101/133520

**Authors:** Micah P. Stumme-Diers, Siddhartha Banerjee, Mohtadin Hashemi, Zhiqiang Sun, Yuri L. Lyubchenko

## Abstract

In the absence of a functioning centromere, chromosome segregation becomes aberrant, leading to an increased rate of aneuploidy. The highly specific recognition of centromeres by kinetochores suggests that specific structural characteristics define this region, however, the structural details and mechanism underlying this recognition remains a matter of intense investigation. To address this, High speed atomic force microscopy was used for direct visualization of the spontaneous dynamics of CENP-A nucleosomes at the sub-second time scale. We report that CENP-A nucleosomes change conformation spontaneously and reversibly, utilizing two major pathways: unwrapping, and looping of the DNA; enabling core transfer between neighboring DNA substrates. Along with these nucleosome dynamics we observed that CENP-A stabilizes the histone core against dissociating to histone subunits, unique from H3 cores which are only capable of such plasticity in the presence of remodeling factors. These findings have implications for the dynamics and integrity of nucleosomes at the centromere.

## INTRODUCTION

The centromere is a specialized locus possessed by all chromosomes which aids in the segregation of sister chromatids during cell division. Failure to faithfully segregate chromosomes will result in aneuploidy, a hallmark of cancer (1). Furthermore, the presence of more than one centromere can lead to di-centric attachments, which can further result in chromosome breakage (2,3). A matter of outstanding importance is to investigate specific features of centromere derived nucleosomes which might directly aid in its role as a substrate for microtubule attachment. Numerous studies over the past two decades have shown that the characteristic feature of centromeres is the presence of centromere protein A (CENP-A); a histone in nucleosomes that replaces the H3 histone of bulk chromatin. Therefore, it is widely accepted that CENP-A nucleosomes epigenetically define the centromere and form the foundation on which the kinetochore is built.

Structural characterization of CENP-A nucleosomes by X-ray crystallographic studies revealed that the exit/entry DNA segments of the nucleosomes (13 bp) are detached from the histone core, thus shortening the DNA segment wrapped around the core by 26 bp when compared to those that contain H3 (4,5). This deficit in DNA wrapped around CENP-A nucleosomes is further supported by enzymatic digestion experiments (6). Hence, one might ask how these differences in nucleosomal DNA lengths can be translated to the structural uniqueness of CENP-A containing nucleosomes *in vitro* and to what degree the replacement of H3 with CENP-A affects spontaneous nucleosome dynamics.

Several recent structural studies have demonstrated the degree to which H3 and CENP-A containing histone cores can tolerate deformation. Biophysical pulling assays revealed that CENP-A is slightly more amenable to disruption than corresponding H3 nucleosomes (7); a finding that agrees well with recent all-atom molecular dynamics (MD) simulation and FRET studies (8,9). Furthermore, using high-resolution NMR spectroscopy, remodeling factors were shown to increase the plasticity of H3 histone cores, indicating that they are capable of far greater flexibility than previously supposed (10).

We recently proposed a model that explains how the shortening of nucleosomal DNA can affect the structural properties of nucleosomes, their arrays, and eventually higher order chromatin structures that define the non-canonical properties of the centromere (11). Furthermore, we hypothesize that the flexible CENP-A nucleosomes are more dynamic, which contributes to the overall structural and dynamic properties required for a functioning centromere (8,12).

In this study, we performed direct imaging of CENP-A nucleosomes by high-speed time lapse atomic force microscopy (AFM), enabling us to visualize dynamics of the particles at a sub-second frame rate (11,13,14). Nucleosomes used for evaluation of DNA wrapping around the histone core were assembled on a DNA substrate containing a centrally positioned 601 motif (14-17). The broadly dynamic behavior of the DNA flanks was first revealed by analysis of AFM images acquired in ambient conditions. Time-lapse imaging further identified dynamic pathways unique to CENP-A nucleosomes that were not previously observed for H3. The spontaneous unwrapping of DNA flanks can be accompanied by the reversible and dynamic formation of loops with sizes equivalent to a single wrap of DNA. Furthermore, CENP-A nucleosome cores are capable of reversible translocation over the DNA substrate; a process mediated through the formation of internal DNA loops along the nucleosome core particle. Finally, the transfer of the histone core from one DNA substrate to another was visualized. The data demonstrate that CENP-A nucleosomes are very dynamic, permitting it to distort freely and reversibly, which may play a critical role in centromere integrity during mitosis and replication.

## MATERIAL AND METHODS

### Preparation of DNA Substrate and Nucleosome Reconstitution

The DNA substrate used in nucleosome assembly contained the 147 bp 601 strong positioning sequence (18) flanked by plasmid DNA, 154 and 122 bp in length, generated using PCR with the pGEM3Z-601 plasmid (Addgene #26656; Fig. S1). Nucleosome assembly was achieved using the salt gradient dialysis method (19,20). Briefly, in 20 µL of 1X TE buffer containing 2M NaCl, recombinant human CENP-A/H4 tetramer’s and H2A/H2B dimers purchased from EpiCypher (#16-0010 and #15-0311, respectively; Research Triangle Park, NC) were mixed with purified DNA substrate (final concentration of 100 ng/uL) at a DNA/dimer/tetramer molar ratio of 1:2.2:1. Each assembly was loaded onto a Slide-A-Lyzer MINI dialysis unit (3,500 MWCO, Thermo Fisher Scientific) which was next dialyzed at 4°C while exchanging the 2M NaCl dialyzate from with one containing 2.5 mM NaCl (both at pH 7.5) at a rate of ∼0.5 ml/min. After 70 hours, the reaction mixture was dialyzed against fresh low salt (2.5 mM NaCl) buffer for another hour. The reassembled nucleosome stock solution was stored at 4°C in the final dialysis buffer (2.5 mM NaCl, 1X TE, pH 7.5). The H3 octamer, CENP-A assembly mixture, and assembled CENP-A nucleosomes were run on a 12% SDS-PAGE gel (Fig. S2). H3 containing octamers were purchased from EpiCypher (#16-0001) and were assembled on the 601-substrate using the method just described.

### Atomic Force Microscopy (ambient conditions)

A 167 µM solution of 1-(3-aminopropyl)-silatrane (APS) was used to modify freshly cleaved mica for 30 minutes at RT as previously described (21,22). The nucleosome stock solution was diluted to 2 nM (based on DNA concentration) in a buffer containing 10 mM HEPES (pH 7.5) and 4 mM MgCl_2_. Immediately following nucleosome dilution, 7 µL of sample was deposited on chilled (4°C) APS-mica and was rinsed with 1.5 mL of ultrapure H_2_O after a two-minute incubation followed by drying with a light flow of argon air. Prepared samples were stored under vacuum until imaged on a Multimode AFM/Nanoscope IIId system using TESPA probes (Bruker Nano Inc). A typical image captured was 1.5 × 1.5 µm in size with 512 pixels/line.

### High-Speed Atomic Force Microscopy

A piece of mica, ∼100 µm thick, was punched into a circle of 2 mm diameter and glued on the stage of the HS-AFM where it was functionalized with 2.5 µL of a 500 µM solution of APS for 30 minutes followed by a rinse with 20 µL of ultrapure H_2_O. Nucleosome stock was diluted to 2 nM in buffer containing 10 mM HEPES (pH 7.5) and 4 mM MgCl_2_ (imaging buffer) and 2.5 µL of the diluted sample was deposited onto the APS-mica for 2 minutes followed by multiple rinses with a total of 20 µL of imaging buffer. The sample surface was kept wet through the entire sample preparation process. Images were captured in the imaging buffer by high-speed AFM (RIBM) using an Olympus Micro Cantilever (BL-AC10DS-A2) that was EBD treated. Typical images acquired were 200 × 200 nm in size at a scan rate ranging from 0.2-0.4 sec/frame.

### Analysis and measurement parameters of nucleosomes imaged in air

The samples deposited on APS mica and imaged by AFM in air were analyzed using Femtoscan Online for the length of each free DNA arm. The contour lengths of DNA particles not bound by histone proteins were first measured and a histogram with a bin size of 5 nm produced a single Gaussian fit with a center at 137 nm (SD = 5.5 nm, R2 = 0.99) which was established as the mean value of free DNA contour length. From this fit, a length conversion unit of 0.32 nm/bp was determined and used for subsequent calculations of DNA bp’s free and wrapped around the histone core. For consistent measurements of nucleosome arm lengths, each free DNA arm was measured from the strand end to the core center as shown in Fig. 2*B* and the FWHM was later subtracted from the sum of the free arms to yield the length of unwrapped DNA. The resulting value was subtracted from the statistically determined 137 nm length of the free DNA to obtain the length of wrapped DNA. The length of wrapped DNA was converted to bp using the length conversion unit established above. Gwyddion was used in obtaining height profile curves that were used to determine height, full width at half max height (FWHM), and volume measurements for nucleosome core particles. The three height and FWHM values for each nucleosome were averaged and these values were used in calculation of the nucleosome core particle volume. The averaged curves were plotted as a histogram and fit with a Gaussian curve (Fig. S3).

### Analysis and measurement parameters of nucleosomes imaged by HS-AFM

Movies captured with HS-AFM were initially analyzed using the FalconViewer extension to IgorPro software. Each image was flattened using either plane or line background removal depending on the background quality. Events of interest were saved as tiffs and analyzed further in Femtoscan Online and Gwyddion as done for the dry images. A moving median was plotted along with raw data for visualization of the global dynamics taking place. Images published had an additional band pass filter applied to lower noise; this filter was not applied to images used in analysis. OriginPro 2016 was used for all graphs both in text and supplement.

## RESULTS

### DNA wrapping of CENP-A nucleosomes characterized by AFM

Mononucleosomes containing human CENP-A were reconstituted on a 423 bp DNA substrate containing the 147 bp Widom 601 positioning sequence (18,19) between 122 and 154 bp flanks of non-specific plasmid DNA (Fig. 1*A* and Fig. S1). Static imaging by AFM in ambient conditions revealed centrally positioned nucleosomes, which are distinguishable from ‘naked’ DNA by the wrapped histone core which appears as a bright round feature flanked by two arms of unbound DNA (Fig. 1*B*). The contour lengths of all free DNA molecules were measured and plotted as a histogram (Fig. S3*A*). Fitting a Gaussian curve to the histogram produced an average at 423 ± 17 bp. To determine the length of DNA wrapped in a nucleosome, the contour length of each free DNA arm was measured and subtracted from the 423 bp length of the ‘naked’ DNA. Each measurement was made from the end of the strand to the center of the core for consistent measurement and average full width at half max (FWHM) of three cross section height profiles of each core particle was subtracted from the sum of its respective arms (Fig. 1*B*). A histogram of the wrapped DNA yielded a bimodal Gaussian distribution with the first peak centered at 120.6 ± 20 bp and a second peak with a center at 160.6 ± 13 bp which were determined to make up 66% and 34% of the total population, respectively (Fig. 1*D*). The population with a peak centered at 120.6 ± 20 bp agrees with the value previously reported (5). A histogram of the heights of nucleosomes produced a bimodal Gaussian distribution with peaks centered at 2.1 ± 0.16 nm and 2.5 ± 0.17 nm which were calculated to populate 36% and 64% of total nucleosomes, respectively (Fig. 1*E*). Similar results were obtained for the nucleosome volumes (Fig. S3*D*). These results suggest that taller particles correspond to the population of nucleosomes with 120.6 bp wrapped DNA. A close inspection of nucleosomes with 160.6 bp of wrapped DNA revealed a heterogeneity in nucleosome core morphology. Strikingly, an unexpected observation was that a significant population of such CENP-A nucleosomes contain DNA looped out of the particle (Fig. 1*F*). As a control, mononucleosomes containing H3 were assembled on the 601 substrate and were prepared for imaging under the same conditions as CENP-A (Fig S4*A*). The DNA wrapped by these particles was determined to be 143.9 ± 20 bp which is in line with the expected value of 147 bp (Fig S4*B*). The contrast in dynamics between CENP-A and H3 nucleosomes, including the looping of the former, is further established below.

**Figure 1.**
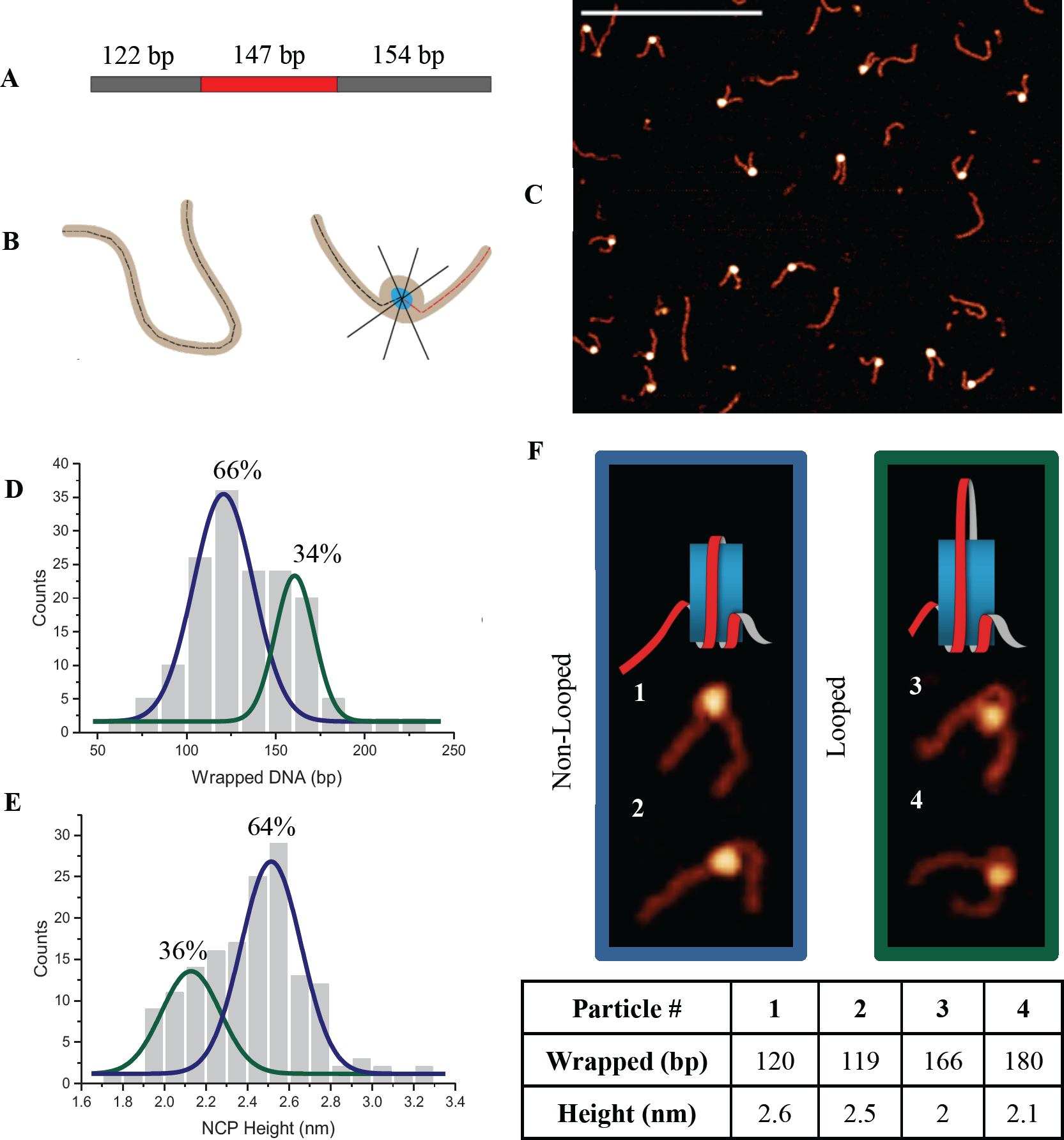
Ambient imaging of CENP-A nucleosomes reveals presence of looped nucleosome structure. (*A*) Schematic of the DNA substrate containing the 147 bp 601 positioning sequence (red) with flanking 122 and 154 bp plasmid DNA (grey). (*B*) Diagram of how (left) the contour length of free DNA was measured and how (right) each nucleosome ‘arm’ was measured from the end of the DNA to the center of the wrapped core (black and red dashed); the width of the particle was then measured by three height profile cross sections at FWHM (black lines through core). Half of this value was subtracted from each arm. (*C*) AFM image of reconstituted CENP-A nucleosomes on an APS-mica surface (scale bar = 400 nm); each round white feature is a histone core with two DNA ‘arms’. Histograms fit with Gaussian curve of: (*D*) bp wrapped DNA with peaks centered at 120.6 ± 20 bp (blue) and 160.6 ± 13 bp (green; bin size = 15 bp, R^2^ = 0.99, 158 counts total). (*E*) nucleosome height with peaks centered 2.1 ± 0.16 nm (green) and 2.5 ± 0.17 nm (blue; bin size = 0.1 nm, R^2^ = 0.95, 158 counts total); Blue curves correspond to non-looped nucleosome and green curves to looped. Percentages above each plot represent the contribution of that conformation to the total nucleosome population based on the integral of the curve. (*F*) Nucleosomes selected from each population seen in the height and wrapped DNA histograms. The color of each box represents the population shown by that color curve in (*D* and *E*). The height and wrapped DNA values for each of these particles is in the table below. The cartoon above the images is a visual of the non-looped and looped nucleosome structure.

### Dynamics of CENP-A nucleosomes visualized by time-lapse AFM

The rather broad distributions of all parameters (wrapped DNA, height, volume) of CENP-A nucleosomes on the 601 substrate obtained at ambient conditions suggests a highly dynamic behavior relative to H3. To directly characterize such dynamic events, we turned to time-lapse AFM, in which nucleosome samples are imaged in aqueous solution. Specifically, we used high-speed AFM which allows for direct observation of the nucleosomes at a sub-second image acquisition rate (13,14,23). Fig. 2*A* shows a few frames out of the 632 consecutive frames assembled as Movie S1. The selected area contained four CENP-A nucleosomes; the dynamics of which were captured simultaneously at a rate of 3.3 frames per second.

**Figure 2.**
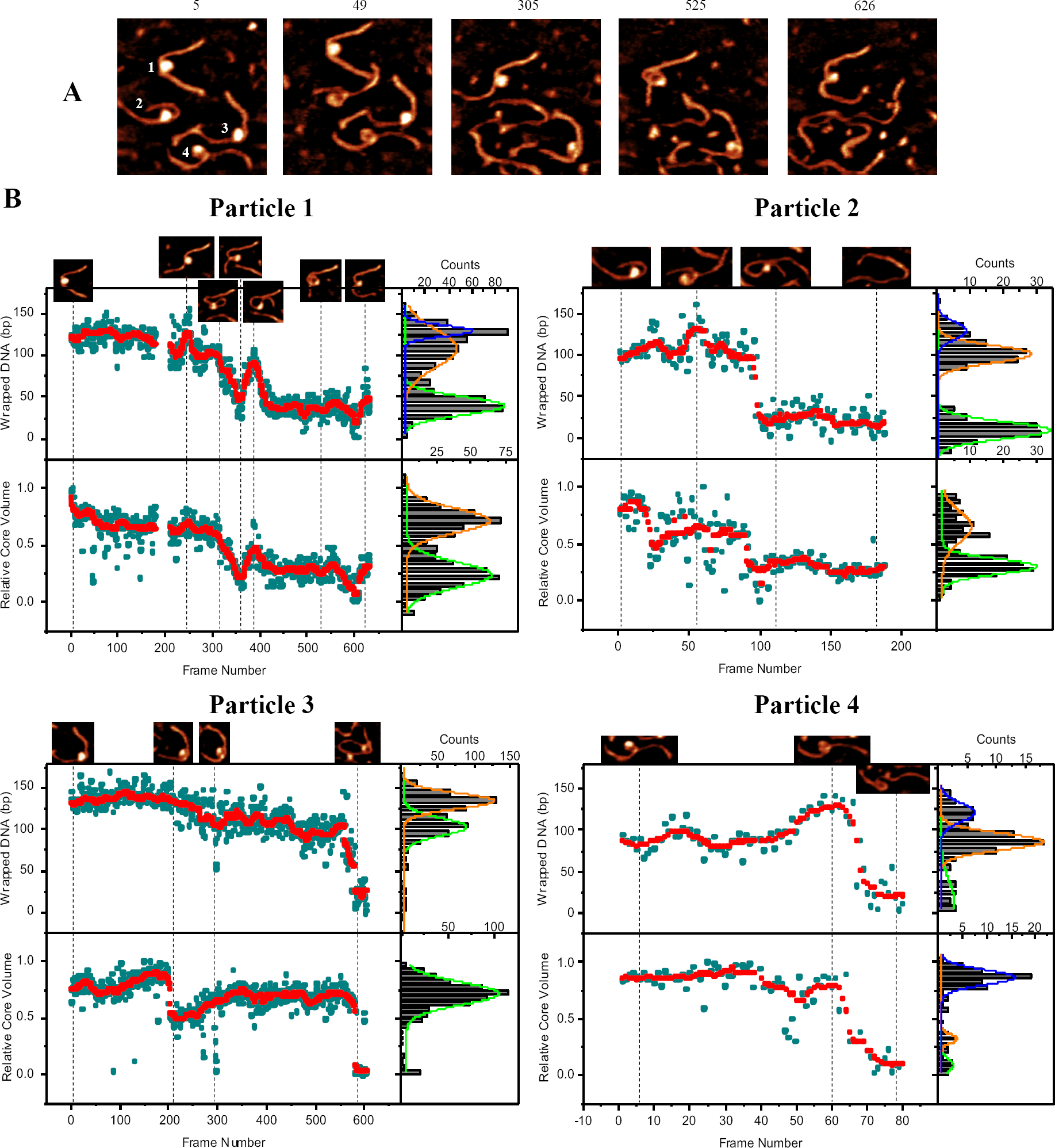
Visualization of CENP-A nucleosomes by HS-AFM reveals increased dynamic behaviors. (*A*) HS-AFM images representing “snapshots” of the diverse dynamics of four nucleosomes across 632 consecutive frames captured at a a rate of 3.3 frames/sec as can be seen in Movie S1. The frame number corresponding to the time at which each frame was captured is above each frame and the four particles are referenced according to their assigned number in the first image. Each image is 200nm × 200 nm. (*B*) Analysis of the wrapped DNA (bp) and relative core volume (normalized 0 to 1) as a function of time. Raw data is shown as cyan circles. Moving median (i±5; red) of the raw data makes the overall dynamic trends clear. The raw data from each particle was plotted as a histogram and fit by a Gaussian curve with centers at the following values where error is represented by std. dev. of the curve: Particle 1; wrapped DNA = 125 ± 15, 105 ± 26, 38 ± 12 bp (R^2^ = 0.99), volume = 0.67 ± 0.1, 0.29 ± 0.1(R^2^ = 0.95), each histogram total count = 598. Particle 2; wrapped DNA = 126 ± 15, 100 ± 11.4, 21 ±12.5 bp (R^2^ = 0.98), volume = 0.64 ± 0.15, 0.30 ± 0.08 (R^2^ = 0.90), each histogram total count = 188. Particle 3; wrapped DNA = 134 ± 11, 103 ± 12.5 bp (R^2^ = 0.99), volume = 0.72 ± 0.11 (R^2^ = 0.96), each histogram total count = 605. Particle 4; wrapped DNA = 121.4 ± 11, 86.7 ± 13, 17.1 ± 15 bp (R^2^ = 0.99), volume = 0.87 ± 0.06, 0.32 ± 0.04, 0.09 ± 0.06 (R^2^ = 0.88), each histogram total count = 79. Bin size for wrapped DNA = 10 bp and for relative core volume = 0.05. Total measurements made for analysis of these particles exceeds 15,000.

Every nucleosome in this dataset spontaneously unwraps and the dynamics were characterized by measuring the lengths of DNA arms. These measurements were made for each of the four particles across all 632 frames as presented in Fig. 2*B* and Fig. S5-7. The two shortest-lived particles, particles 2 and 4, are initially wrapped less than the 121 bp mean value (Fig. 1*D*), with 100 ± 11 and 86 ± 13 bp, respectively. Next, the partially unwrapped nucleosomes begin a wrapping process, achieving wrapped values of 126 ± 15 bp and 121 ± 11 bp for particles 2 and 4, respectively. The transient wrapping is followed by unwrapping of both particles until they become fully unwrapped after ∼100 frames (Fig. 2*B*; particles 2 and 4).

Particles 1 and 3 (Fig. 2*B*) demonstrate a dynamic behavior different from that of 2 and 4. Initially both nucleosome cores are wrapped to values close to the expected 121 bp and DNA dissociates from each of the cores gradually over the ∼600 frames. Particle 1 begins with 125 ± 15 bp of wrapped DNA for ∼200 frames until it unwraps to 105 ± 26 bp for ∼200 frames. This is a process with multiples unwinding-rewinding step as is evident from the large width of the Gaussians approximated from the experimental histograms (see Fig. 2*B*). Unique from the other four nucleosomes, particle 1 never fully unwraps, instead rewrapping 38 ± 12 bp around a smaller histone complex, where it remains stable for over 200 frames. Particle 3 begins wrapped with 134 ± 11 bp, where it remains for over 400 frames, until partially unwrapping ∼30 bp of DNA to 103 ± 12.5 bp, where it remains until it eventually fully unwraps (Fig. 2*B*).

### Dynamic formation of DNA loops

As discussed above, AFM images of dry nucleosome samples with looped out nucleosomal DNA were identified (Fig. 1*F*). Given that looping out of DNA in chromatin is an issue of great biological importance (24-26), we next focused our attention on dynamics of this specific behavior. A few frames representing snapshots of this looping process were selected from Movie S2 and are shown in Fig. 3*A*. These images visually illustrate that the loop grows gradually (frames 1 through 3), until it reaches a size of ∼90 bp, as seen in frame 4. Next, the loop shrinks (frames 5 though 7) becoming again small (∼20 bp) in frame 8. To characterize this process, images of all 100 frames were analyzed. It was done by contour length measurements of the looped DNA, height profile analysis of a line dissecting the apex of the loop and core, relative volume of the nucleosome core particle, and by changes in the DNA wrapped around the core via nucleosome arm contour measurements. These are plotted in Fig. 3*B-E*. Contour length analysis of the loop (Fig. 3*B* and *C*) shows that its formation is reversible, as it dramatically changes size five times as seen on the loop cross sections and the loop size graph. The first two times the loop is formed, it grows to be ∼15 bp until retightening around the core. Shortly after, the formation of the first of these two loops was found to be accompanied by a more than two-fold reduction in relative core volume where it remained for the subsequent looping events (Fig. 3*C* and *E*). This reduction in volume can be attributed to an approximately two-fold reduction in relative height of the nucleosome, while the width of the particle is seen to slightly increase, as also revealed by still images. Such a reduction in height can be explained in part to a loss of DNA from the histone core due to looping, which greatly affects such measurements. The potential dissociation of H2A/H2B was not observed during the loop formation events. A height reduction of this magnitude which is never reached again despite shrinking of the loop, suggests that upon looping of the DNA, an internal rearrangement of the histone core has occurred. Unlike the first two loops formed, the third grows to a size of ∼30 bp where it remains stable for eight frames until growing to ∼90 bp. After five frames, the ∼90 bp loop shrinks back to ∼30 bp where it remains stable for over 20 frames.

**Figure 3.**
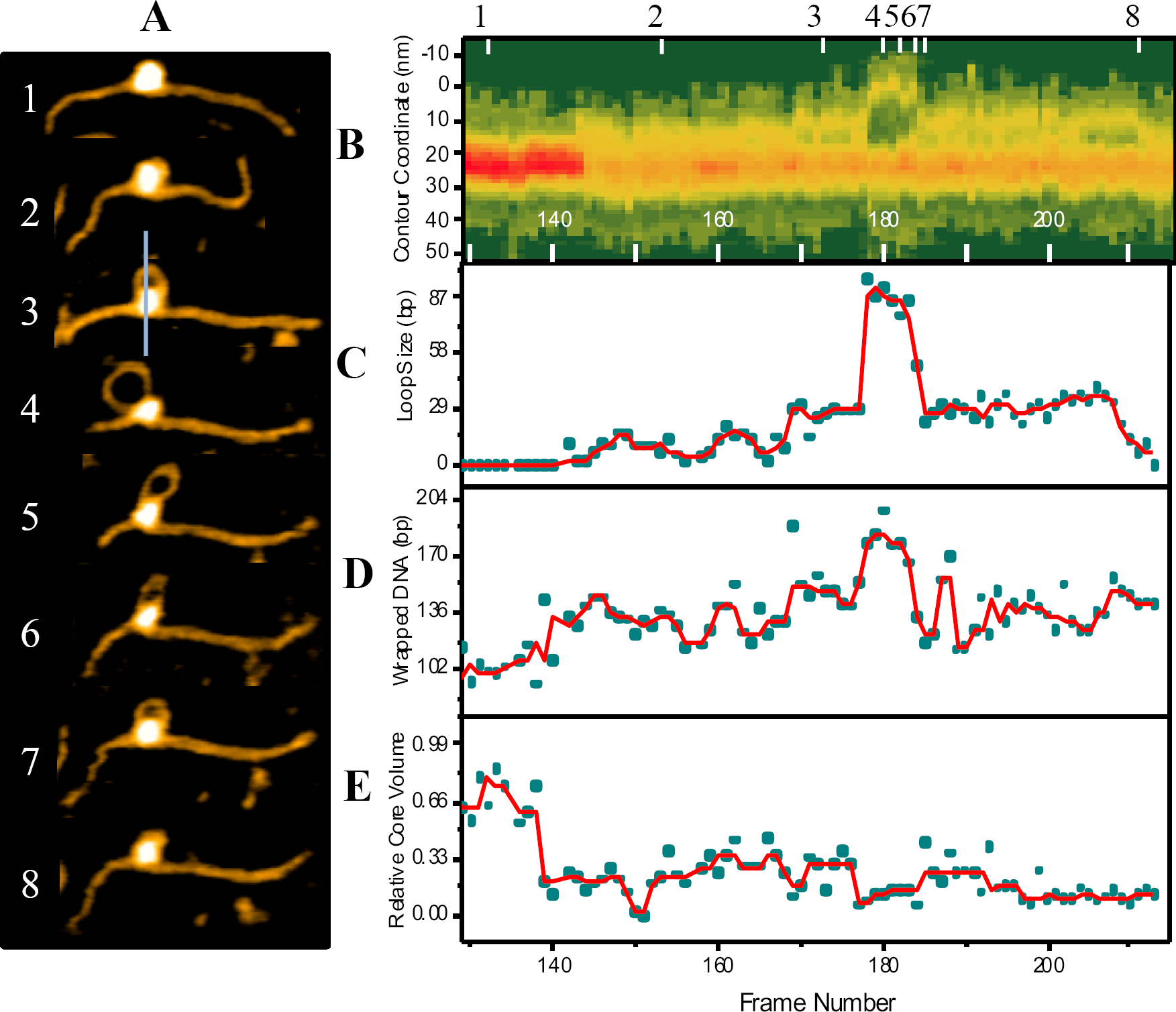
Spontaneous looping of CENP-A nucleosomes visualized by high speed AFM. (*A*) Images representing stages of the looping of DNA from the nucleosome complex were selected from Movie S2; numbers (1-8) correspond to the location of the frames (*A*) in the time trajectory. The line in image three shows how height profile cross sections were measured to obtain plot (*B*); this represents the cross sections as a function of frame number where the maximum height of the core is aligned for each cross section. Deviations in the upper yellow line make clear the dynamic nature of this looped complex. The reduction in height shown as a transition from red to yellow immediately prior to loop formation at approx. frame 140 suggests a rearrangement of the histone core as the DNA loosens around it for loop formation (*C*) A plot of loop size as a function of time shows the dynamic growth and shrinking of the loop. (*D*) Wrapped DNA (bp) was determined by arm length measurements which revealed that changes in arm do contribute to the formation of the loop. (*E*) The core volume was found to decrease by two times initially which is largely due to a reduction in height. Raw data is represented as cyan circles, and the moving median by a red line. In total, looping was observed for a total of 10 nucleosomes from the 52 particles imaged.

Measurements of the DNA arm contour lengths lend insight to the mechanism by which the loop is forming; either by sliding of the DNA around the particle, or by falling away of the DNA from the core particle. As each of these loops forms, one of the DNA arms remains relatively stable, while the other is seen to decrease in size in unison with loop growth, suggesting that the loop is formed through the sliding of one of the DNA arms around the histone core. Since these arm lengths are used in the calculation of the wrapped DNA, a decrease in the length of the arm produces an artificially high value for bp of wrapped DNA, as was observed in AFM images in ambient conditions (Fig. 1*D* and *F*).

### Translocation and transfer of nucleosomes

Previous studies have illustrated that the spontaneous unwrapping of nucleosomes is the major DNA dissociation event for H3 containing chromatin subunits; as was also observed for H3 nucleosomes assembled in this study (Movie S3 and Fig. S8). was also observed However, potential translocation (or, hopping), disruption of protein-protein contacts, or sliding of the histone core were not taken into consideration. In our previous time-lapse experiments we did visualize sliding, although it was limited to a relatively short range (14). Long-range sliding was only previously observed in the presence of the detergent, 3-[(3-Cholamidopropyl)dimethylammonio]-l-propanesulfonate (CHAPS), suggesting that such long range translocation of canonical nucleosomes is only possible in the presence of CHAPS (14,16). In contrast, CENP-A nucleosomes behave rather differently and are capable of long-range translocation. One of such events is shown in Fig. 4; this set of images (out of >500 assembled as Movie’s S4 and S5) demonstrates that translocation takes place over 180 bp and that the process is bi-directional. The initial translocation takes place while the nucleosome is partly unwrapped, and the DNA moves in a corkscrew like motion around the core until it is at the end of the DNA substrate (Fig. 4*A* and Movie S4). The contour length of the ‘shrinking’ DNA arm was measured from its end, to the center of the core particle for 250 frames until the core had moved to the end of the DNA substrate. This ‘forward’ translocation moves the DNA a total of ∼180 bp but then appears to stall twice along the way. The first time is after the DNA moves ∼25 bp and the second after it moves ∼75 bp. Lastly, it moves ∼70 bp to the end of the DNA strand where it stays for few hundred frames (Fig. 4*A* and *B*). Next, the DNA begins a reverse translocation of ∼180 bp, back to its starting position (Fig. 4*B* and Movie S5). Unlike the partly wrapped, corkscrew-like translocation, the reverse propagation stalls four times after moving the following distances in order of the translocation: ∼60 bp, ∼52 bp, ∼40 bp, ∼25 bp (Fig. 4*B*). Similar translocation events were clearly observed for seven nucleosome particles.

**Figure 4.**
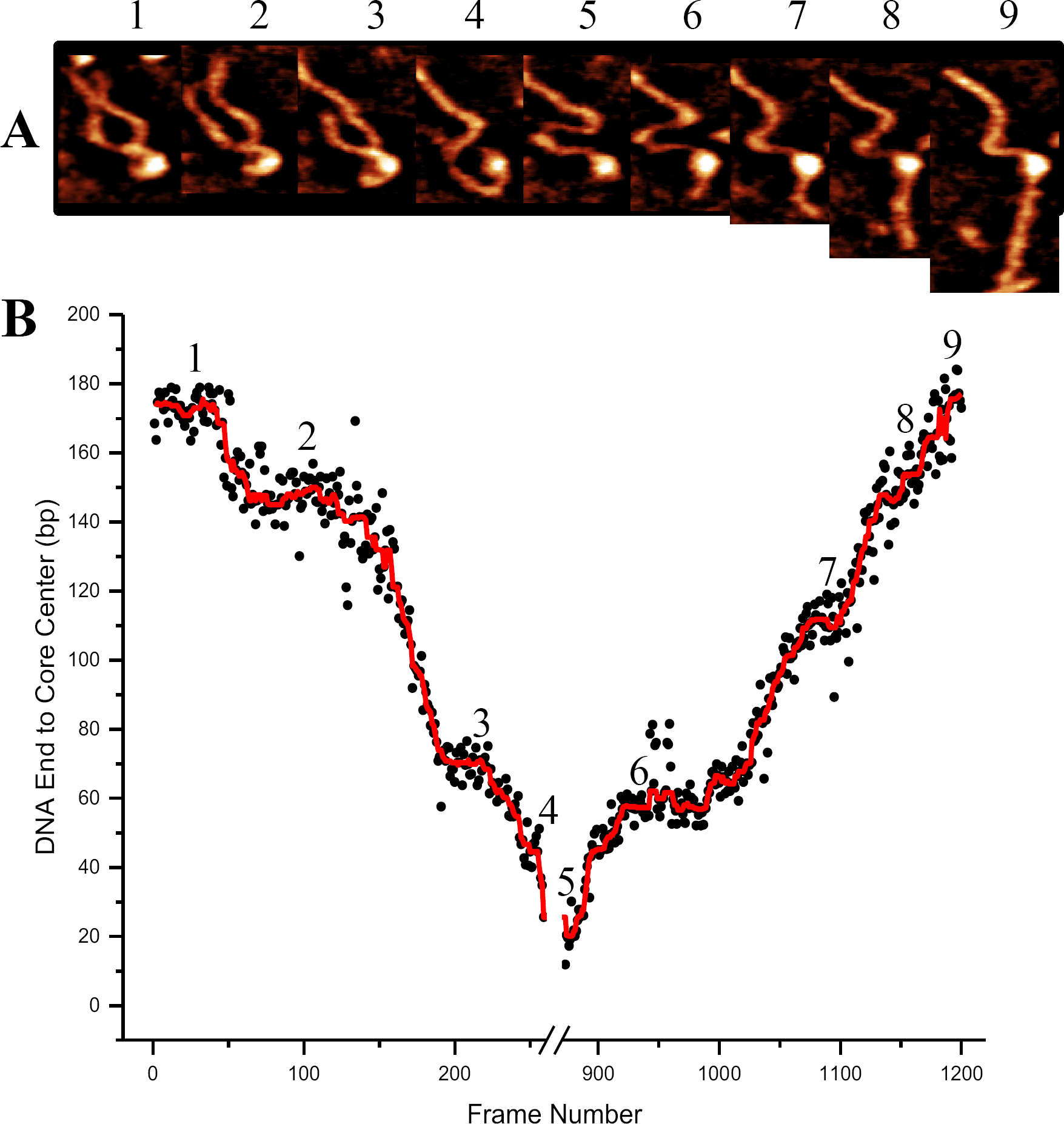
Nucleosome translocation is reversible along a DNA substrate. (*A*) Images of the forward (1-4) and reverse (5-9) translocation of a nucleosome core particle. Forward movement of the complex appears to be achieved via a corkscrew motion of the DNA as best visualized in Movie S4. The reverse movement is of a less wrapped complex and ends up at about the same position on the DNA substrate as where it began (Movie S5). The white arrows in images 1 and 6 point to the arm length for which the contour length measurements are shown in (*B*) where the distance was measured from the end of that arm to the center of the core particle for every frame of the video. Frames between ∼250 and ∼850 were excluded because the core remained at the end of the DNA substrate (as seen in image 5) for this time and no translocation took place. Black circles represent the data points and the red line is a moving median. Sliding was observed for a total of 6 particles from the 52 particles imaged.

Following the reverse translocation, the core particle was seen to transfer from one DNA molecule to another. A set of a few AFM images is shown as Fig. 5*A*; the full set of data can be seen in Movie S6. Initially, two nucleosomes are shown in the same scanning area. One nucleosome unwraps leaving a free DNA substrate (frame 2) which is later approached by the second nucleosome core particle (frame 3). This nucleosome core particle becomes associated simultaneously with two DNA molecules (frame 3 and 4), eventually transferring its core to the free DNA substrate. Schematically this process is shown in Fig. 5*B*.

**Figure 5.**
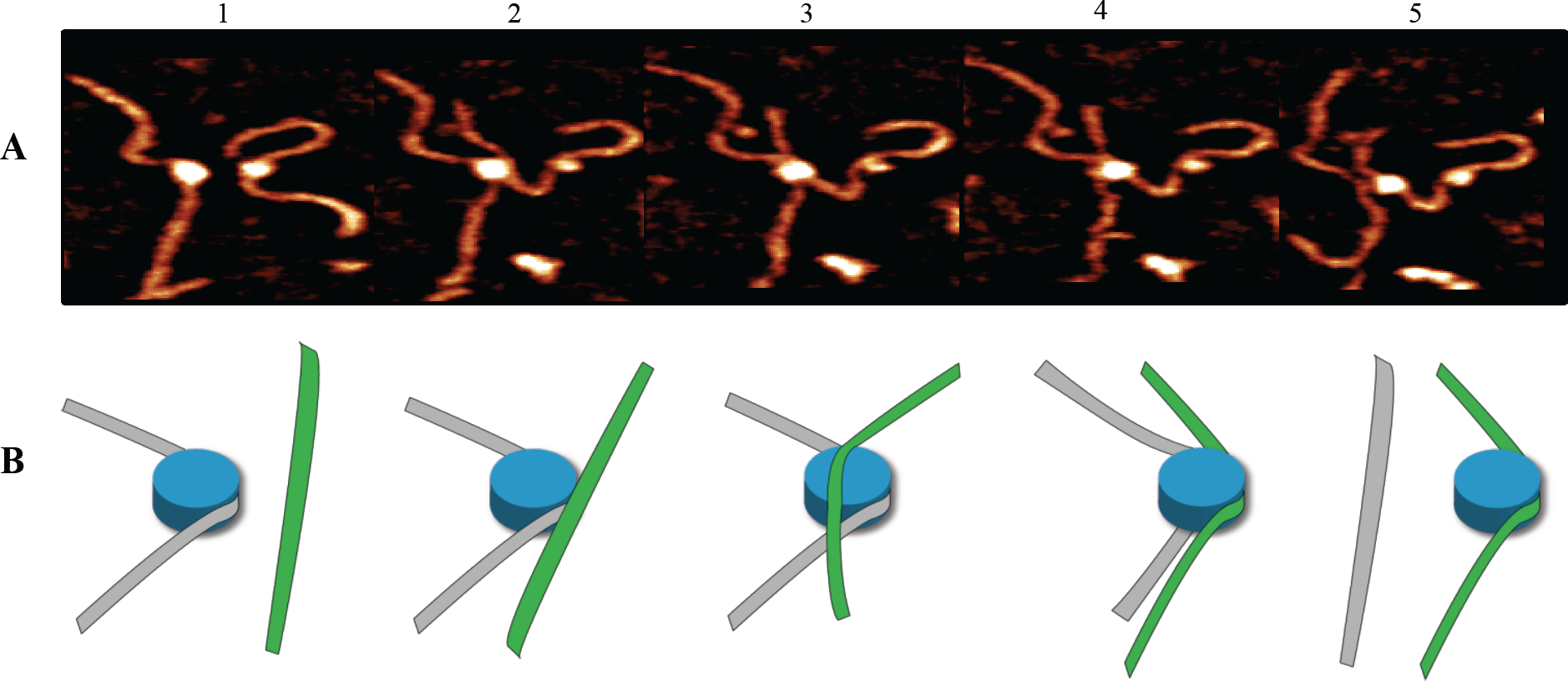
The stable CENP-A histone core is capable of spontaneous inter-strand transfer. (*A*) HS-AFM images depicting the transfer of a partially wrapped CENP-A containing histone core from one strand to another as seen in Movie S6 and depicted in schematic (*B*). Image one shows the parent strand (grey) on the left is partly wrapped with the histone core (blue) free from interaction with the acceptor strand (green) on the right (image and schematic 1). Interaction of the acceptor strand (image and schematic 2) soon leads to a histone/DNA complex containing both parent and acceptor strands (image and schematic 3). Within a second of this dual substrate complex, the acceptor strand took full control of the core with the parent strand only partly interacting (image and schematic 4) until the acceptor/core complex is completely free from interaction with the parent substrate (image and schematic 5).

## DISCUSSION

In this study, we identified the dynamics of CENP-A nucleosomes by AFM techniques including high-speed time lapse imaging which revealed that CENP-A nucleosomes are capable of spontaneous unwrapping. As was reported previously for H3 nucleosomes and observed for the H3 control sample in this study, the spontaneous unwrapping of CENP-A nucleosomes was the eventual fate for all particles imaged (14-16). The simultaneous visualization of four nucleosomes as they unwrap (Fig. 2 and Movie S5) illustrates that the phenomenon is common and that spontaneous dynamics are an intrinsic feature of all nucleosomes. Unlike H3 containing nucleosomes however, the unwrapping process/pathway for each CENP-A nucleosome is different; some of them unwrap rapidly like H3 (e.g., particles 2 and 4), while others take a longer time via dynamic pathways (e.g., particles 1 and 3). While a similar process was observed for canonical nucleosomes, we identified a number of dynamic behaviors unique to CENP-A nucleosome that are not found in the canonical nucleosome (14,17). Details of these features are discussed below.

### Nucleosome unwrapping via loop formation

A distinct feature of CENP-A nucleosomes is the bimodal distribution of wrapped DNA as shown in Fig. 1*F*. The major peak is associated with wrapping ∼120 bp DNA which makes ∼1.5 turns around the CENP-A core compared with ∼1.7 turns for canonical particles. This value is in line with recent crystallographic and Cryo EM studies (5,27). This difference was discussed and highlighted in our recent paper in which different models of nucleosomal arrays based on these differences in turn number were proposed, as supported by several studies (5,11,27). However, we identified a second population of nucleosomes with much shorter DNA flanks that were determined to have 160 bp of wrapped DNA which corresponds to as many as two full turns of DNA around the core. This population was found to contain approximately half as many particles as the main population and is unique to CENP-A nucleosomes, as such a bimodal distribution is not observed with canonical nucleosomes which have been shown both in this (Fig S4*B*) and previous studies to have a single population with ∼1.7 DNA turns (15,28). High-resolution images in Fig. 1*F* revealed that the arm length deficit for a portion of the CENP-A population is not due to the elevated wrapping of DNA but rather to large segments of DNA that are looped out from the core. These looped nucleosomes are formed in such a way that the non-looped DNA, in contact with the core, consists of both 601 sequence and an adjacent segment of non-specific DNA. The mechanism by which such loops assemble was next probed using time-lapse imaging in which (Fig. 3) we observed small loops are formed initially which continue to grow over time. Two processes contribute to the looping-out of DNA: First, a segment of DNA dissociates from the histone core, as evidenced by decrease the particle size (compare frames 1 and 4 in Fig. 3*A*). Second, a segment of the left flank moves around the core. Importantly, the segment remained bound to the histone core forming a stably existing loop. As it is seen from frames 5 through 8, the loop shrinks and this process is primarily due to moving of the left flank of the DNA substrate. The looping process is reversible due to the retained integrity of the CENP-A histone core.

### Translocation of histone core

The spontaneous translocation of CENP-A histone cores is another dynamic property of CENP-A nucleosomes as illustrated in Fig. 4. In this example, the core moves over ∼180 bp and then eventually returns to the initial position. It stops at the end of the DNA substrate and has a few pauses along the translocation path. A similar pattern is observed for the reverse translocation. We used arm length measurements to estimate the DNA wrapping efficiency of the histone core during the translocation. Initially (frame 1 in Fig. 4*A*.), 104 bp of DNA are seen to wrap around the histone core, which is 16 bp shorter than for the mean value for CENP-A nucleosomes estimated in Fig. 1*D*. However, only a portion (∼40 bp) of the DNA is in direct contact with the core, as the rest forms a looped structure. As translocation progresses, this loop structure changes in size, which results in fluctuations of the DNA segment in contact with the core. Once at the end of the DNA substrate, the loop unwraps, leaving ∼40 bp of DNA still bound to the core. Spontaneously the distal end of DNA moved beyond the scan size, so determination of nucleosome size was further made through volume measurements. The results shown in Fig. S9 demonstrate that the nucleosome volume fluctuates, but generally remains constant, suggesting that during translocation the CENP-A core remains associated with ∼40 bp. This is confirmed when the distal DNA end later reenters the frame and arm measurements provided the expected DNA contact value of ∼40 bp.

Translocation has previously been observed for H3 nucleosomes and is therefore not unique to CENP-A (14). However, a distinction can be made when comparing the ranges of H3 translocation to that of CENP-A. H3 is limited to ∼40 bp as the histone core quickly loses integrity and dissociates from DNA into its histone components (14). Ranges exceeding 100 bp were only achieved in the presence of CHAPS detergent, which like other detergents stabilizes the histone core from dissociation in dilute solutions of mononucleosomes (14,16). In this study, we revealed that CENP-A is capable of reversible long range translocation of ∼180 bp in the absence of stabilizing factors/conditions. Critical to this ability is the stability of the CENP-A histone core upon loosening or unwrapping of DNA. Despite losing two-thirds of its contacts with DNA prior to translocation, the core remains whole and continues spontaneous interactions with and along the DNA substrate. As CENP-C has been shown to stabilize fully wrapped CENP-A nucleosomes, our results suggest that the unwrapped core is also stable and capable of maintaining dynamic contacts with DNA (9).

### Interstrand transfer of CENP-A core

Transfer of the nucleosome core from one DNA substrate to another, as illustrated in Fig. 5, is another characteristic of CENP-A nucleosomes that has not been directly visualized before. Evidence of interstrand nucleosome transfer was presented and discussed in landmark studies (29,30) and was recently demonstrated in a magnetic tweezers study (31). Furthermore, our results directly support such an ability. An interesting feature of the transfer process, as Fig. 5 illustrates, is that the transfer takes place between two DNA substrates. Note that exchange between the two DNA strands takes place with the CENP-A nucleosome core in contact with ∼40 bp, as schematically shown in the cartoon (Fig. 5*B*). We would like to emphasize two important features of the CENP-A nucleosome core that makes such a transfer possible: the retained integrity of the histone core, and the long lifetime of the core-DNA complex after unwrapping. Again, these properties are unique to the stable CENP-A core and were not previously observed for canonical nucleosomes (14).

Overall, the experiments presented here reveal the highly dynamic properties of CENP-A nucleosomes that contribute to centromeric chromatin remodeling. Nucleosomes spontaneously undergo the unfolding process utilizing two major pathways: looping, and translocation along DNA, which enables the transfer of nucleosomes from one DNA to another. These spontaneous processes can be modulated by environmental conditions. Nucleosome remodeling can occur by different pathways with the necessary pathway for a specific genetic process (repair, replication, transcription) being promoted through recruitment of a specialized remodeling factor (32); for example, Chd1 can be used for sliding of nucleosomes (33). Our discoveries of the nanoscale dynamics intrinsic to CENP-A nucleosomes suggest that dynamic rearrangements of centromeric chromatin can occur in the absence of remodeling factors and at the same time facilitate dynamics of chromatin catalyzed by the remodeling factors. In addition to CENP-A nucleosome dynamics, we observed that CENP-A stabilizes nucleosome core particles against complete dissociation upon loosening or unwrapping of DNA. We speculate that this novel property of CENP-A might permit it to stay associated with H2A, H2B and H4, thereby permitting rapid nucleosome re-assembly following mitosis, transcription or replication induced eviction. Furthermore, the stabilization of fully wrapped CENP-A nucleosomes by CENP-C (9) along with the intrinsic stability of unwrapped CENP-A cores, may contribute to its longevity on the chromatin fiber, thus contributing to the cell cycle independent “memory” of centromeric chromatin over several cell cycles (34).

## ACKNOWLEDGEMENT

We thank Yamini Dalal for the gift of recombinant histones, training, enthusiastic feedback, and critical review of the manuscript. We acknowledge Daniel Melters (Dalal lab) for thoughtful discussions and critical review of the manuscript. Author contributions: YLL and MSD designed the project; MSD, SB, MH, ZS performed AFM experiments and data analyses. All authors wrote and edited the manuscript.

## FIGURES

Begin on the following page (13) in the order that they appear.

## SUPPLEMENTARY DATA

Supplementary figures 1-9 can be found in the attached PDF labelled “Supplementary_Data”. Movies 1-6 are attached as.mov files (Movie S1-S6) and their descriptions follow:

**Movie S1.** Relative dynamic behavior of four CENP-A nucleosomes imaged simultaneously. This demonstrates the consistent fate of all nucleosomes imaged with other dynamic events taking place prior to spontaneous unwrapping. Captured at a rate of 3.3 frames per second for a total of 632 frames. Analysis of the dynamics of each particle are shown as Fig. 2 and Fig. S5-7.

**Movie S2.** Looping of DNA from CENP-A nucleosome. Prior to unwrapping of DNA from the histone core, a loop of DNA can spontaneously grow from the wrapped nucleosome and can then tighten back around the histone core. Captured at a rate of 2.7 frames per second for a total of 107 frames. Analysis of looping dynamics are shown as Fig. 3.

**Movie S3.** Spontaneous unwrapping of H3 nucleosome on 601 substrate. In line with previous studies, the H3 nucleosome spontaneously unwraps. Captured at a rate of 2.1 frames per second for a total of 167 frames. Analysis of unwrapping is shown as Fig. S4*B*.

**Movie S4.** Forward long range translocation of CENP-A histone core along DNA substrate. The core moves from the substrate center to the end until again translocating back to the center (Movie S5). Captured at a rate of 3.3 frames per second for a total of 284 frames. Further analysis is presented as Fig. 4.

**Movie S5.** Reverse long range Translocation of CENP-A histone core along DNA substrate. The same particle as shown in Movie S4 moves back to the position where it began in the middle of the substrate. Between the forward translocation in Movie S4 and this reverse translocation, the core remained at the very end of the DNA substrate (as it begins in this reverse translocation movie) and were therefor excluded from the movie. Captured at a rate of 3.3 frames per second for a total of 328 frames. Further analysis is presented as Fig. 4.

**Movie S6.** Interstrand transfer of CENP-A histone core. Following the forward and reverse translocation (Movie S4 and Movie S5, respectively) the CENP-A core transfers to an adjacent DNA substrate. Captured at a rate of 3.3 frames per second for a total of 290 frames. Still frames of the transfer and a cartoon depicting this process are shown as Fig. 5*A* and *B*, respectively.

